# Sex-specific growth in invasive pink salmon (*Oncorhynchus gorbuscha*) from two regions in northern Norway

**DOI:** 10.64898/2026.11.02.763571

**Authors:** Erlend Aaberge Nes, August Gjestvang, Kathrine Dunlop, Christian Selbach, Eva B. Thorstad, Kenyon B. Mobley

## Abstract

The recent invasion of pink salmon (*Oncorhynchus gorbuscha*) along the Norwegian coast has raised significant ecological concerns. Investigating regional differences in marine growth is an important step in evaluating the establishment of pink salmon in new environments. Pink salmon were captured in two northern Norwegian fjords during their return migration to in the summer of 2023; Varangerfjord in Finnmark and Oksfjord in Troms, representing the eastern and western extent of the core areas of the pink salmon invasion. Body size estimates (total length, weight, condition factor) were used to identify differences in sex-specific marine growth at the time of capture. Scales were analyzed to estimate marine somatic growth via the distance between circuli and the position of the last winter circuli (LWC). Estimates of cumulative circuli distance were divided into four marine growth periods that roughly correspond to 1) spring/summer, year one, 2) summer/fall, year one, 3) fall/winter, year one, and 4) spring, year two. Sex-specific and location effects were found in the body size of pink salmon, where males were heavier and had higher body condition than females and Varanger males had the highest estimates of body size. Scale readings revealed that growth rates were highest in the spring of both years (periods 1 and 4), with significantly decreased growth in the fall/winter (periods 2 and 3). Males from the established Varanger location exhibited the highest growth (17-21% total cumulative circuli distance) over all periods compared to the other groups, suggesting that these fish experience more favorable survivorship and marine growth conditions. These insights into pink salmon marine growth emphasize the importance of taking sex and location into consideration in further assessing the species’ expansive potential in Norway.

## Introduction

Understanding differences in traits such as growth, behavior, and reproduction along an invasion front is an important step in evaluating invasive species success in new environments. Individuals at the leading edge of an invasion often exhibit traits that enhance dispersal and population expansion compared to those in established populations (Sakai et al., 2001). These individuals at the invasion front may grow faster, reproduce earlier, or display behaviors that promote movement into new areas, a pattern driven by processes such as spatial sorting and rapid evolutionary change (Burton et al., 2010; Perkins et al., 2013). For example, studies have shown that invasion-front populations can evolve increased dispersal ability and altered life-history traits, accelerating the spread of the species and intensifying ecological impacts (Phillips & Shine, 2006). Recognizing these differences may help to improve predictions of invasion dynamics and inform targeted management strategies (Shine, 2012).

In this study, we investigate sex-specific somatic growth of the invasive pink salmon, *Oncorhynchus gorbuscha*, from two geographic locations in Northern Norway. The pink salmon is an anadromous species of salmon native to both sides of the northern Pacific Ocean (Heard, 1991; Sandlund et al., 2019). Pink salmon were introduced to the Barents Sea after the USSR began stocking rivers in 1956 with fry originating from the eastern Pacific Ocean (Gordeeva et al., 2015). Introduction efforts persisted until 2000, after which, every following generation originated from natural reproduction (Erkinaro et al., 2026). While pink salmon have sporadically been observed in Norway since the early 1960’s, the numbers have been generally low and mostly contained north of 70° latitude (Erkinaro et al., 2026). In 2017, a massive increase in abundance and southern expansion was observed, with around 6000 pink salmon registered in at least 272 Norwegian rivers (Mo et al., 2018). By 2023, pink salmon abundance has increased 50-fold in Norway compared to abundances recorded in 2017, with the majority of fish in northern Norwegian Rivers (Staveley et al., 2025). For example, pink salmon abundance in the Tana River have increased by 3500% since 2017 (Erkinaro et al., 2026) illustrating the urgent need to address pink salmon invasion and its ecological impacts.

Pink salmon possess numerous traits contributing to its successful colonization (Pess et al., 2012). Among these traits is a pattern of a higher straying rate in their homing behavior compared to other salmonids (Hendry et al., 2003). While the extent of this rate is debated and dependent on various factors such as sex, stream type, location, and hatchery produced versus wild fish stock (Mortensen et al., 2002), studies from Alaska have reported straying rates over 50% in streams more than 40 km from hatchery release sites (Brenner et al., 2012). In comparison, straying rates observed in the Varzuga River in the White Sea exceeded 15% over distances of 10 to 70 km (Sandlund et al., 2019). Pink salmon have also demonstrated a high capacity for rapid adaptation, likely due to strong selection and high generation turnover (Kovach et al., 2012). This adaptability is demonstrated by the populations of freshwater exclusive pink salmon, established unintentionally in the Great Lakes, USA in the 1950’s (Kwain & Rose, 1986; Langan et al., 2026). The capacity for adaption and expansion emphasize the challenge of managing further westward and southward expansions of pink salmon in Norway.

Studies conducted in pink salmon’s native range in the northeastern Pacific Ocean have found survival is dependent on marine growth (Moss et al., 2005), which is largely influenced by ecological conditions such as sea temperature and diet (Beamish et al., 1999; Hindar et al., 2020; Mortensen et al., 2000). As a result, survival rates of pink salmon vary across geographic regions (Pyper et al., 2001). Studies from pink salmon’s native range have found faster growth in the early marine phase (sea entry until early fall) significantly affects survival rate of individuals over the following winter indicating that this is a critical phase in the pink salmon life cycle (Moss et al., 2005). With survival and return rates of mature pink salmon affecting their spread and establishment, growth rate may provide an indirect measure of how successfully the species is adapting to local environmental conditions.

As pink salmon are a relatively newly established species, the marine migration and feeding patterns is not well known in Norway (Diaz Pauli et al., 2022) although our knowledge is growing. For example, Paulsen et al. (2022) found evidence indicating that Norwegian pink salmon experience a critical period of marine growth before winter, while also establishing that growth conditions vary between years. Diaz Pauli et al. (2022) reported a trend of body size increases towards higher latitudes. Additionally, Diaz Pauli et al. (2022) observed a differing prey composition in pink salmon stomachs between northern and southern marine feeding grounds. Variability in marine growth was also found in pink salmon sampled between years and two regions in Norway (Simmons et al., 2026). Mid and late season growth during the first year was also found to have a strong influence on adult size at the time of capture (Simmons et al., 2026). Oceanic conditions may also influence abundance and distribution of pink salmon in Norway, with a particular concern that warming oceans could provide more favorable conditions to this species (Hindar et al., 2020).

Sexual differences in growth are likely important during the onset of maturation after the first winter at sea in pink salmon. During maturation, males typically grow larger and develop pronounced secondary sexual traits such as humped backs and elongated jaws or kypes, which enhance success in competition for mates (Beacham et al., 1988; Heard, 1991; Quinn, 2018). Female pink salmon typically mature at smaller sizes and invest less into secondary sexual traits (Beacham et al., 1988; Heard, 1991; Quinn, 2018). Instead, females prioritize energy investment into gonad development and egg production (Dickerson et al., 2002; Gende et al., 2004; Iritani et al., 2025). These differences influence survival, reproductive output, and population dynamics, as males prioritize mating competition and females optimize fecundity (Dickerson et al., 2002; Heard, 1991). Environmental conditions such as temperature (Jeffries et al., 2011) and food availability (Lennox et al., 2023) may further amplify or constrain these sex-specific growth patterns, making sexual dimorphism an important consideration for marine growth, but remains poorly understood in the invasive range.

In this study, body size data, and scale samples collected from pink salmon in Norway were analyzed to investigate regional and sex-specific differences in marine growth and size at capture. Analyzing fish scales is a widely used method for studying somatic growth in pink salmon (Kaev, 2015; Kaev et al., 2023; Moss et al., 2005; Paulsen et al., 2022). These analyses are based on measuring distances between circuli, or microstructures deposited in the scales that reflect somatic growth in pacific salmonids (Courtney et al., 2000; Fukuwaka, 1998; Fukuwaka & Kaeriyama, 1997; Kaev, 2015). These measurements can thereby be used to assess growth patterns over time and spatial scales and are highly correlated with body length at capture in Pacific salmonids (Courtney et al., 2000; Paulsen et al., 2022; Simmons et al., 2026; Walker & Sutton, 2016).

The study targeted two locations in northern Norway, Varangerfjord located in Finnmark within the eastern core area of the pink salmon invasion, and Oksfjord, Troms representing the western invasion front. Pink salmon were collected during their second summer at sea in coastal marine waters at the onset of sexual maturation and the return migration. We compared the total length, weight and the condition factor between males and females at the two locations to examine sex- and region-specific differences at the time of collection. Somatic growth was measured by scale analysis over four distinct time periods 1) spring/summer, year one, 2) summer/fall, year one, 3) fall/winter, year one, and 4) spring, year two. We hypothesized that potential differences in somatic growth between regions along the invasion front that might reflect differences in survivorship and environmental conditions experienced by fishes inhabiting these two regions. Additionally, we hypothesized that sex-specific growth would occur in the second year at the onset of maturity. Potential differences in sex-specific growth between regions may indicate changes in growth, maturation and reproduction that may inform invasion dynamics and the potential for successful establishment of the species.

## Methods

### Study species

Pink salmon have a two-year semelparous lifecycle, with a high level of genetic divergence between odd and even year-classes (Beacham et al., 1985; Christensen et al., 2021; Hawkins et al., 2002). Pink salmon in Norway appear to be from odd year stock, with high abundances recorded in odd years (Staveley et al., 2025; Thorstad et al., 2026). Spawning migration in Norway begins in mid-June and continues into August (Thorstad et al., 2025). Spawning in Finnmark typically takes place between early August to mid-September (Lennox et al., 2023; Thorstad et al., 2026). Hatching usually occurs during winter or spring (Sandlund et al., 2019), depending on the timing of spawning and location as embryo development is highly temperature dependent (Dunlop et al., 2026; Lennox et al., 2023). Between March and May, pink salmon fry emerge from the riverbed, measuring about 30 mm in length (Sandlund et al., 2019). At this point, fry are already capable of hypo-osmoregulation (Gallagher et al., 2013) and are believed to migrate towards seawater shortly after (Sandlund et al., 2019). In Finnmark, the timing of seaward migration appears to vary both within and between rivers, with observations spanning from mid-May (Sandlund et al., 2019), to late June (Bjerknes, 1977; Erkinaro et al., 2024). After a year of marine growth, mature adults return to freshwater to spawn, and subsequently die shortly thereafter (Sandlund et al., 2019).

### Sampling

Pink salmon were collected from two location in northern Norway (Fig. 1) during two cruises organized by the Norwegian Institute of Marine Research (IMR) in the summer of 2023. On June 27, 2023, adult fish were obtained from Varangerfjord in Finnmark County, 69°54′25.20″N, 21°15′55.70″E using a bag net deployed from the RV Fangst as part of a study to develop capture methods for pink salmon on the coast while reducing bycatch of other salmonids. The bag net is a passive capture method with a leading net that guides fish into a capture chamber. The sampling site was located near Storsand at the southern mouth of the Varangerfjorden, a fjord located on the southern flank of the Varanger peninsula. Fish were collected as bycatch between July 9 and July 14, 2023, from Oksfjord in Troms County, Norway, 70°01’07.5”N, 29°25’21.4”E in fyke-nets used to collect anadromous Arctic charr (*Salvelinus alpinus*) as part of the national salmon lice monitoring program (Norwegian Institute of Marine Research). Oksfjord is an inlet on the eastern side of Reisafjorden in Nordreisa municipality, approximately 14 km^2^ in size.

**Figure 1.**
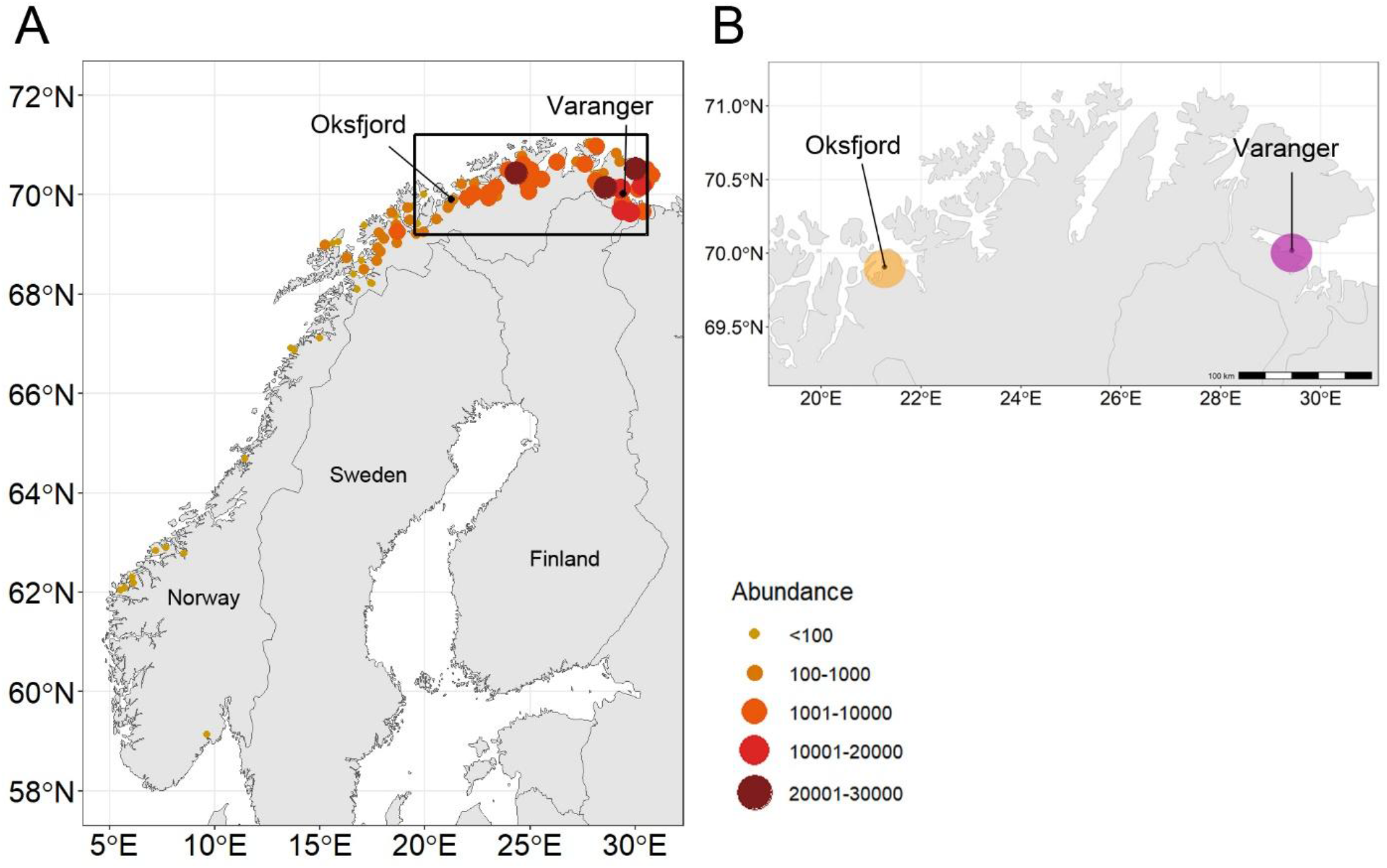
A) The distribution and abundance of pink salmon in 2023. Data from Miljø-direktoratet (2026) based on landings from targeted fishing measures in 93 rivers. B) Inlay box shows the position of the pink salmon collection locations, Oksfjord, Troms County (orange) and Varangerfjord (Varanger), Finnmark County, Norway (Purple).

All fish were immediately frozen after capture and the transported to the Norwegian University of the Arctic, Tromsø (UiT) for further analysis. Fork length (mm) and wet weight (g) were measured, and sex was determined by inspecting gonads after dissection for each fish. Condition factor (K) was calculated using Fulton’s condition factor formula, where W is weight (g), and L is the fork length (cm) (Heincke, 1908):

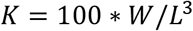

Using a scalpel (size 24), mucus was removed, and scales scraped off in the area posterior to the dorsal fin and dorsal to the lateral line. The scales were stored in paper envelopes and stored at room temperature (20-25°C).

### Scale data analysis

Approximately ten scales per fish were selected and imprinted onto plastic slides using a roller press. The resulting plastic impressions were then examined using an Indus 4601-01 desktop reader (Indus International, Wisconsin, USA) to evaluate the readability of the circuli. Scale impressions were then photographed using a Nikon Digital Sight DS-Ri1 (Nikon, Tokyo, Japan) camera attached to a Leica Z6 AP0 macroscope (Leica Geosystems, Wetzlar, Germany) using 32X magnification.

As only a single scale image was used per individual fish for analysis, selection of the scale image was essential to ensure reliable and consistent data collection. The optimal scale images were characterized by clearly defined, unbroken, and non-overlapping circuli, in addition to a distinct focus area. Images containing artifacts such as mucus and foreign particles obscuring the scale circuli, or regenerated scales (i.e., a replacement scale that grows back after a fish loses an original scale due to injury, handling, or abrasion) were also excluded from analysis.

Before counting and measuring circuli, a radius was determined by identifying a straight trajectory line from the center of the focus towards the scale edge (Fig. 2). This line was oriented towards the longest axis of the scale, reflecting the primary direction of growth. Serving as a reference axis, the line was used to count visible circuli starting from the center of the focus, designated as “C0” (Fig. 2). The circulus distance, or the distance between a given circulus (*C*) and its preceding circulus (*C – 1*) was measured along the axis using ImageJ version 1.54g (Schneider et al., 2012). The last winter circulus (LWC) was identified as the final circulus within the band of narrow-spaced circuli before the second summer growth (Fig. 2). The LWC was determined visually and supported by circuli measurements. Total circuli refers to the total number of circuli measured in each scale, LWC distance represents the distance from the center of the focus up to the LWC, and mean circuli distance is the average distance across all measured circuli.

**Figure 2.**
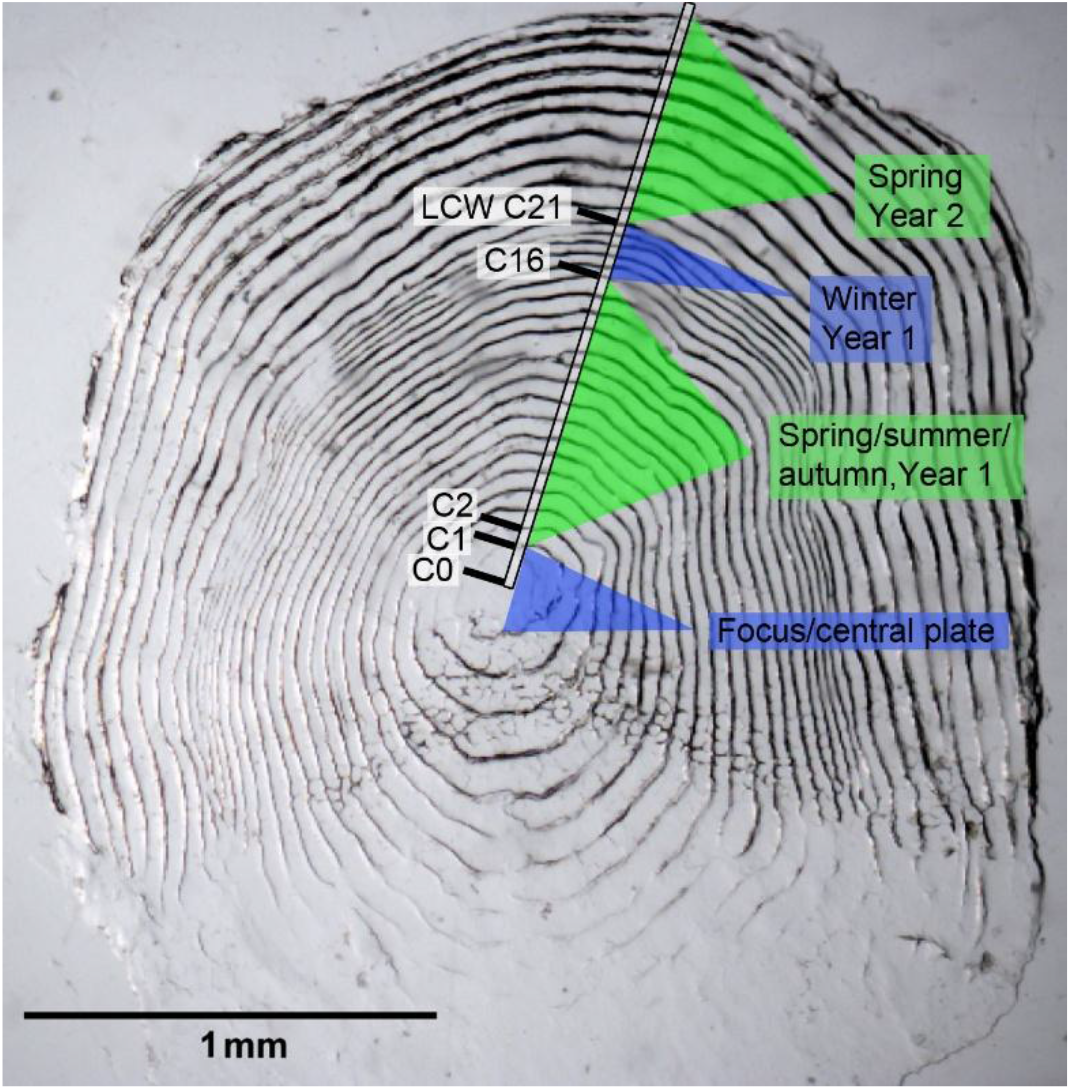
Photo of a pink salmon scale impression showing the radius used to measure circuli distance. The center of the focus area is labeled C0, with circuli marked sequentially. Seasonal zones are identified, including spring/summer/autumn, year one (C1-C16), winter, year one (C16-C21), and spring, year two (>C21). The last circulus in the winter zone (C21) denotes the last winter circulus (LWC).

To assess seasonal growth patterns, the circuli of each scale sample were divided into four growth periods based on the position of the LWC, functioning as a reference point for end of winter. The first three periods divided the circuli pre LWC into equal thirds. When the number of pre-LWC circuli was not evenly divisible by three, the remaining number of circuli was distributed between the two first periods. While this approach does not account for individual variation in circuli deposition, the growth periods based on the common winter circulus reference (LWC) should correspond to roughly the same time periods across individuals. Period four was defined as the six circuli following the LWC, resetting the LWC as 0 so that the following six circuli represented the initial growth rate in the spring of the second year. The periods roughly correspond to 1) spring/summer, year one 2) summer/fall, year one, 3) fall/winter, year one and 4) spring, year two.

### Statistical analyses

To investigate the influence of sex and location on the physical characteristics of pink salmon at the time of capture, linear models (LMs) were used. Fork-length, weight, and condition factor representing body size metrics were analyzed as response variables, with location, sex and their interaction included as explanatory variables. Linear models were also used to uncover variation in scale characteristics. Separate LMs were constructed using sex, location and their interaction as explanatory variables with the total number of circuli and the circuli number of the LWC position as response variables.

The four growth periods were used to investigate if somatic growth, as measured by cumulative circuli distance, differed across developmental stages based on sex or location. First, we investigated how total cumulative circuli distance within each growth period differs with respect to sex, location and their interactions using a linear model approach. Circuli distances within each period were cumulatively added, starting from the first circuli measurement within each given period. This model compares the total amount of growth between periods in relation to sex and location. Second, we investigated the rate of growth within each period with the relationship between the cumulative circuli distances and the circuli number. Because individuals varied in the number of circuli present within each growth period, the circuli number was normalized to have a mean of zero and a standard deviation of one prior to analyses. These cumulative circuli distances were tested within each period using general linear mixed models (GLMMs) with circuli number, sex, location, and their interaction as predictors and the unique ID number as a random factor using the *lme4* version 2.0-1 package (Bates *et al*., 2015) and lmerTest version 3.2-1 (Kuznetsova et al., 2017).

All statistical analyses were performed in R (R Core Team, 2026), at a significance level of α = 0.05. Means are reported as ± 1 standard error of the mean (SE). Models were initially fitted with the full interaction terms of the explanatory variable. The highest order interactions were first tested and removed if found not to be significant (p > 0.05), simplifying the model until only significant interactions or main effects remained. Model assumptions were assessed by simulating residuals using the DHARMa version 0.4.7 (Hartig, 2026) package in R. This assessment involved evaluating residual normality, homoscedasticity, outliers, and uniformity, with visual inspection of Q-Q plots.

## Results

A total of 60 pink salmon were collected; 30 from Oksfjord and 30 from Varangerfjord. Means and sample size for body length, body weight, and condition factor pooled by sex and location are reported in Table S1. The sexual distribution of fish caught in Oksfjord was near unity, with 53% females, while more females were collected in Varangerfjord (63% female).

### Body size at capture

At the time of capture, Varanger males had, on average, longer fork length than Varanger females and Oksfjord males and females (Table S1, Fig. 3). However, fork length was not significantly influenced by sex or location (Sex: 1.2961 ± 1.0004, *t*= 1.295, *p* = 0.200; Location: 1.0227 ± 1.9954, *t* = 1.027, *p* = 0.309) (Table 1, Fig. 3). Males were significantly heavier than females (Sex: 0.2583 ± 0.0939, *t* = 2.751, *p* = 0.008) but this relationship did not differ between locations (Location: 0.1704 ± 0.0934, *t* = 1.823, *p* = 0.073) (Table 1, Fig. 3). A significant sex-specific difference in condition factor was however found showing that males were in higher condition than females (Sex: 0.1081 ± 0.0378, *t* = 2.865, *p* = 0.006) (Table 1, Fig. 3). Additionally, Varanger fish had a significantly higher condition factor than Oksfjord fish (Location: 0.0815 ± 0.0375, *t* = 2.172, *p* = 0.034) (Table 1, Fig. 3). There were no significant interactions between sex and location in any of the body size models.

**Table 1.** Results of linear models showing the effects of sex and location on fork length (cm), weight (kg), and condition factor (*K*). Reference sex is female, reference location is Oksfjord. Significance for bold values indicates α = 0.05.

| **Response variable** | **Predictor** | **Effect size** | **SE** | **t** | **Pr(>\|t\|)** |
|---|---|---|---|---|---|
| Fork length | Intercept | 44.9021 | 0.8754 | 51.296 | **<0.001** |
|  | Sex | 1.2961 | 1.0004 | 1.295 | 0.200 |
|  | Location | 1.0227 | 0.9954 | 1.027 | 0.309 |
| Weight | Intercept | 1.2917 | 0.0822 | 15.719 | **<0.001** |
|  | Sex | 0.2583 | 0.0940 | 2.751 | **0.008** |
|  | Location | 0.1704 | 0.0934 | 1.823 | 0.073 |
| *K* | Intercept | 1.4195 | 0.0330 | 42.998 | **<0.001** |
|  | Sex | 0.1081 | 0.0377 | 2.865 | **0.005** |
|  | Location | 0.0815 | 0.0375 | 2.172 | **0.034** |

**Figure 3.**
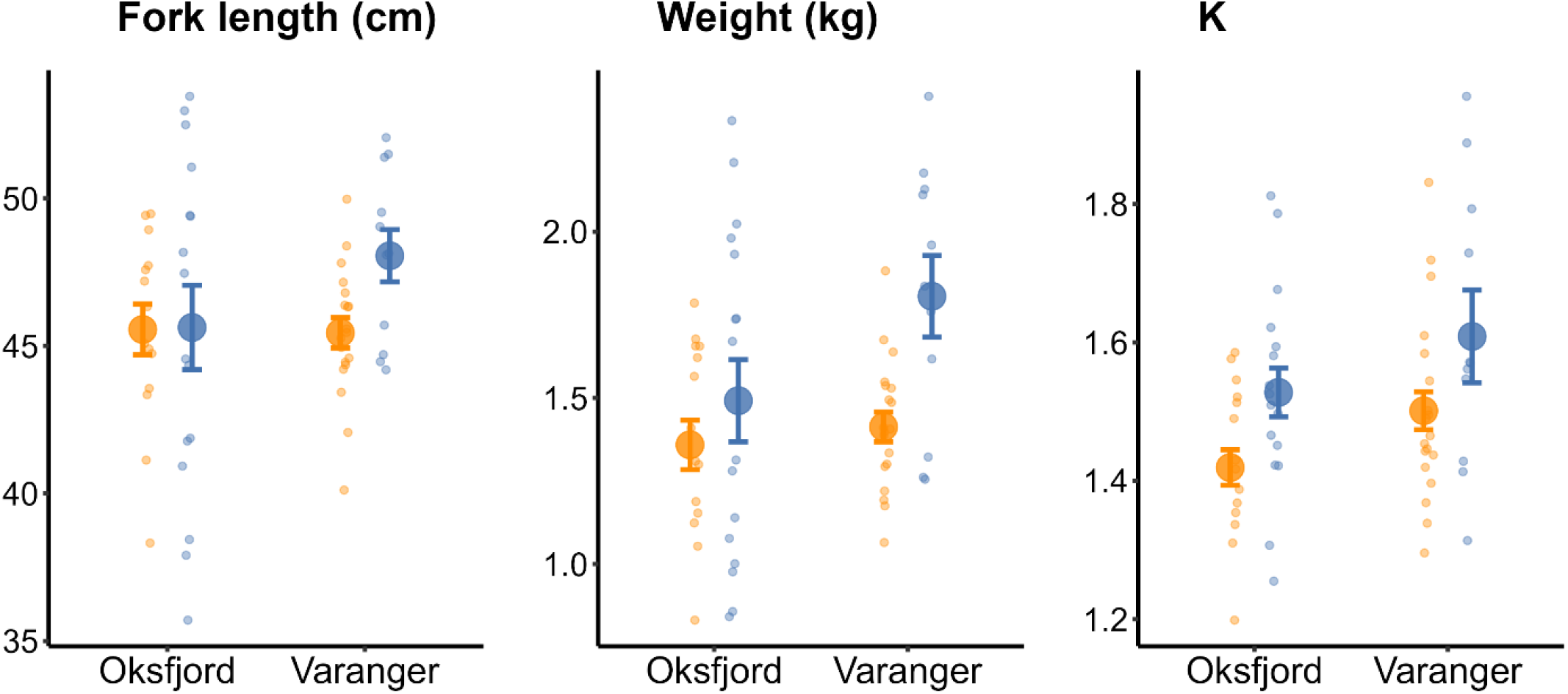
Summary of body size estimates at the time of collection. Mean fork length (cm), weight (kg), and condition factor (*K*) of sampled pink salmon by sampling location (Oksfjord and Varanger) and sex (males in blue, females in orange). Large circles with error bars represent the mean ± SE, while small circles show individual data points. For clarity, individual points are jittered on the x- and y-axis.

### Growth based on scale analyses

Scale measurements from all individuals captured in Oksfjord were analyzed. One male sampled in Varanger was excluded from scale analysis due to having only regenerated scales. Means and sample size for scale measurements including the total number of circuli, position of the last winter circulus counted from the focus, distance from focus to last winter circulus and mean distance between circuli pooled by sex and location are reported in Table S2.

Results of linear models demonstrated that the total number of countable circuli did not significantly differ between the sexes or by location (Sex: 0.3017 ± 0.6381, *t* = 0.473, *p* = 0.638; Location: 0.7006 ± 0.6336, *t* = 1.106, *p* = 0.274) (Table 2). Similarly, the position of the LWC did not differ between sexes or by location (Sex: 0.6507 ± 0.5300, *t* = 1.228, *p* = 0.241; Location: 0.6238 ± 0.5263, *t* = 1.185, *p* = 0.241) (Table 2). There were no significant interactions between sex and location in either the total circuli or position of the LWC models.

**Table 2.** Results of linear models showing the effects of sex and location on the total number of circuli (total circuli) and the position of the last winter circulus counted from focus (LWC position). Reference sex is female, reference location is Oksfjord. Significance for bold values indicates α = 0.05.

| **Response Variable** | **Predictor** | **Effect size** | **SE** | **t** | **Pr(>\|t\|)** |
|---|---|---|---|---|---|
| Total circuli | Intercept | 30.5057 | 0.5532 | 55.143 | **<0.001** |
|  | Sex | 0.3017 | 0.6381 | 0.473 | 0.638 |
|  | Location | 0.7006 | 0.6336 | 1.106 | 0.274 |
| LWC position | Intercept | 19.1863 | 0.4595 | 41.775 | **<0.001** |
|  | Sex | 0.6507 | 0.5300 | 1.228 | 0.225 |
|  | Location | 0.6238 | 0.5263 | 1.185 | 0.241 |

### Growth periods

To evaluate growth over time, total cumulative circuli distance was compared between growth periods, sex and location. We found a significantly lower growth rate in growth periods two and three compared to growth period one and four (Period 2: -0.0381 ± 0.0159, *t* = -2.399, *p* = 0.0173; Period 3: -0.0703 ± 0.0159, *t*= -4.420, *p* < 0.0001; Period 4: 0.0122 ± 0.01590, *t*_12,223_ = -4.420, *p* < 0.0001)(Table 3, Fig 4). Thus, the highest growth rate was encountered in the spring of year one and two, with the lowest growth rates encountered in the summer/fall and fall/winter of year one (Table 3, Fig 4). A significant interaction between sex and location was evident (Sex*Location: 0.05941 ± 0.0130, *t* = 4.571, *p* < 0.0001), indicating significantly higher growth in Varanger males. On average, Varanger males grew 17% more than Varanger females over all periods (Period 1: 16%; Period 2: 23%; Period 3: 18%; Period 4: 10%) based on total cumulative circuli distance. Similarly, Varanger males grew 21% more than Oksfjord males (Period 1: 21%; Period 2: 28%; Period 3: 15%; Period 4: 17%) and 17% more than Oksfjord females (Period 1: 15%; Period 2: 26%; Period 3: 16%; Period 4: 10%)

**Table 3.** Results of linear models showing the effects of sex, location, and growth period on the total cumulative circuli distance. Reference sex is female, reference location is Oksfjord, reference period is 1. Significance for bold values indicates α = 0.05.

| **Predictor** | **Effect size** | **SE** | **t** | **Pr(>\|t\|)** |
|---|---|---|---|---|
| Intercept | 0.0250 | 0.0117 | 21.323 | **<0.0001** |
| Sex | -0.0172 | 0.0143 | -1.203 | 0.2302 |
| Location | 0.0019 | 0.0141 | 0.138 | 0.8907 |
| Period 2 | -0.0381 | 0.0159 | -2.399 | **0.0173** |
| Period 3 | -0.0703 | 0.0159 | -4.420 | **<0.0001** |
| Period 4 | 0.0122 | 0.0159 | 0.767 | 0.4440 |
| Sex:Location | 0.0564 | 0.0130 | 4.571 | **<0.0001** |
| Location:Period 2 | 0.0594 | 0.0182 | 0.814 | 0.4165 |
| Location:Period 3 | 0.0148 | 0.0182 | -0.897 | 0.1706 |
| Location:Period 4 | -0.0163 | 0.0182 | -0.345 | 0.7304 |
| Sex:Period 2 | 0.0130 | 0.0183 | 0.710 | 0.4786 |
| Sex:Period 3 | 0.0032 | 0.0183 | 0.176 | 0.8605 |
| Sex:Period 4 | -0.0077 | 0.0183 | -0.422 | 0.6735 |

**Figure 4.**
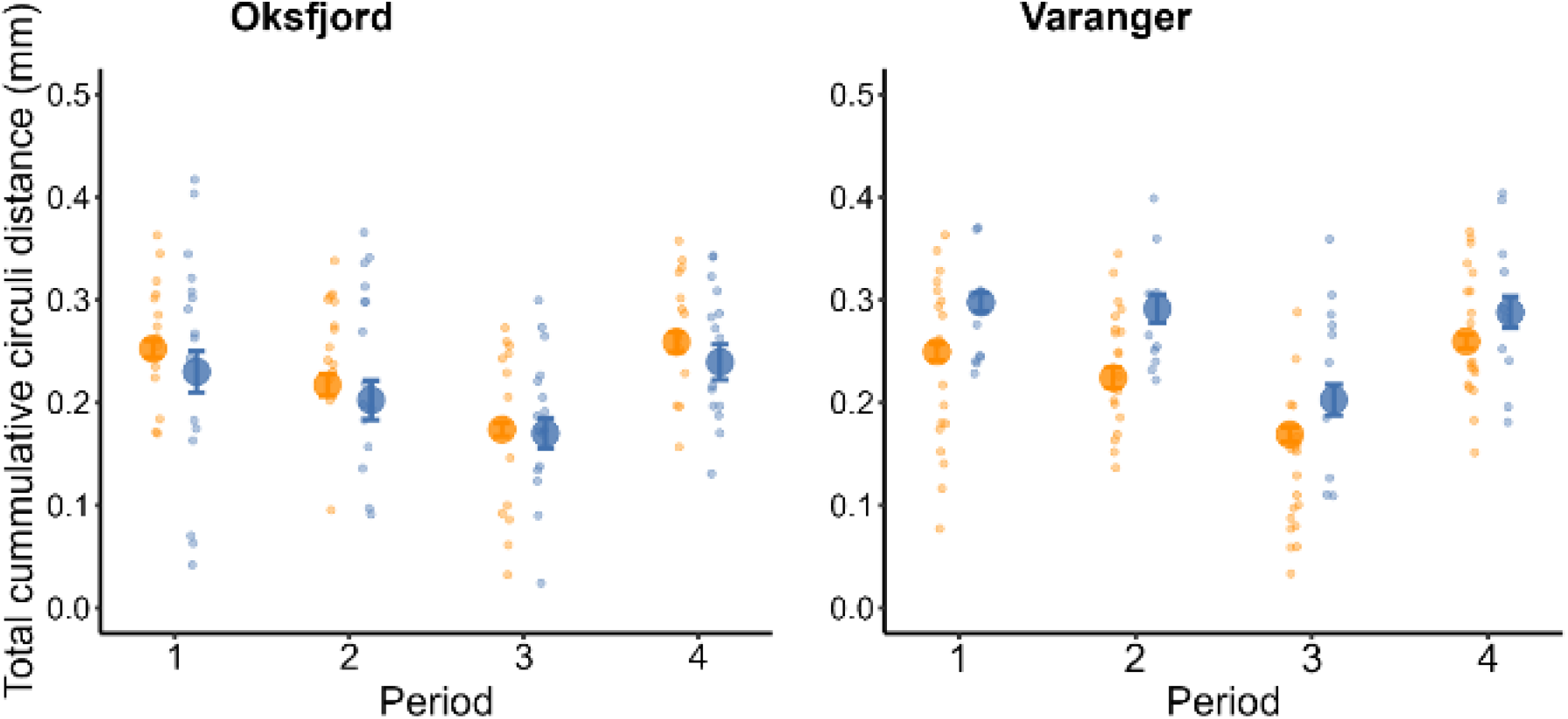
Total cumulative circuli distance (mm) for sampling locations for each sex (males in blue, females in orange) within growth periods. Large circles with error bars represent the mean ± SE, while small circles show individual data points. For clarity, individual points are jittered on the x- and y-axis.

We modeled the growth rate by evaluating the relationship between cumulative circuli distance with circuli number over the growth periods, sex and location using a GLMM approach on normalized circuli number. A significant three-way interaction between circulus number, sex, and location was found in all periods, indicating that the difference in growth rates between sex depended on location, with Varanger males exhibiting faster growth during these times (Circuli no.*Sex*Location: Period 1, 0.0182 ± 0.0036, *t* = 5.031, *p* < 0.0001); Period 2, 0.0021 ± 0.0041, *t* = 5.117, *p* < 0.0001; Period 3, 0.0089 ± 0.0034, *t* = 2.627, *p* = 0.0090; Period 4, 0.0166 ± 0.0042, *t* = 3.986, *p* < 0.0001) (Table 4, Fig. 5). A GLMM on non-normalized raw data also showed similar three-way interactions across all time periods (Table S3, Fig. S1).

**Table 4.** Results of generalized linear mixed models showing the effects of normalized ciruli number (circuli no.), sex and location on cumulative circuli distance (mm) and their interactions for growth periods 1-4. Reference sex is female, reference location is Oksfjord. Significance for bold values indicates α = 0.05.

|  | **Predictor** | **Estimate** | **SE** | **t** | **Pr(>\|t\|)** |
|---|---|---|---|---|---|
| Period 1 | Intercept | 0.1445 | 0.0077 | 18.702 | **<0.0001** |
|  | Circuli no. | 0.0796 | 0.0018 | 43.428 | **<0.0001** |
|  | Sex | -0.0098 | 0.0106 | -0.929 | 0.3567 |
|  | Location | 0.0035 | 0.0102 | 0.347 | 0,7299 |
|  | Circuli no.*Sex | -0.0015 | 0.0025 | -2.321 | **0.0209** |
|  | Circuli no.*Location | -0.0015 | 0.0024 | -0.613 | 0.5402 |
|  | Sex*Location | 0.0509 | 0.0154 | 2.647 | **0.0106** |
|  | Circuli no.*Sex*Location | 0.0182 | 0.0036 | 5.031 | **<0.0001** |
| Period 2 | Intercept | 0.1269 | 0.0081 | 15.730 | **<0.0001** |
|  | Circuli no. | 0.0689 | 0.0021 | 33.116 | **<0.0001** |
|  | Sex | -0.0096 | 0.0111 | -0.871 | 0.3876 |
|  | Location | 0.0017 | 0.0106 | 0.159 | 0.8743 |
|  | Circuli no.*Sex | 0.0003 | 0.0028 | 0.115 | 0.9086 |
|  | Circuli no.*Location | 0.0034 | 0.0027 | 1.245 | 0.2142 |
|  | Sex*Location | 0.0043 | 0.0162 | 2.653 | **0.0104** |
|  | Circuli no.*Sex*Location | 0.0021 | 0.0041 | 5.117 | **<0.0001** |
| Period 3: | Intercept | 0.1074 | 0.0068 | 15.755 | **<0.0001** |
|  | Circuli no. | 0.0547 | 0.0017 | 31.746 | **<0.0001** |
|  | Sex | 0.0010 | 0.0093 | 0.108 | 0.9145 |
|  | Location | -0.0023 | 0.0090 | -0.261 | 0.7952 |
|  | Circuli no.*Sex | -0.0015 | 0.0024 | -0.645 | 0.5195 |
|  | Circuli no.*Location | -0.0014 | 0.0023 | -0.634 | 0.5264 |
|  | Sex*Location | 0.0219 | 0.0137 | 1.606 | 0.1141 |
|  | Circuli no.*Sex*Location | 0.0089 | 0.0034 | 2.627 | **0.0090** |
| Period 4: | Intercept | 0.1345 | 0.0069 | 19.437 | **<0.0001** |
|  | Circuli no | 0.0892 | 0.0021 | 42.920 | **<0.0001** |
|  | Sex | -0.0113 | 0.0095 | -1.194 | 0.2376 |
|  | Location | 0.0007 | 0.0091 | 0.072 | 0.9428 |
|  | Circuli no.*Sex | -0.0070 | 0.0028 | -2.458 | **0.0145** |
|  | Circuli no.*Location | -0.0012 | 0.0027 | -0.428 | 0.6688 |
|  | Sex*Location | 0.0250 | 0.0139 | 1.804 | 0.0767 |
|  | Circuli no.*Sex*Location | 0.0166 | 0.0042 | 3.986 | **<0.0001** |

**Figure 5.**
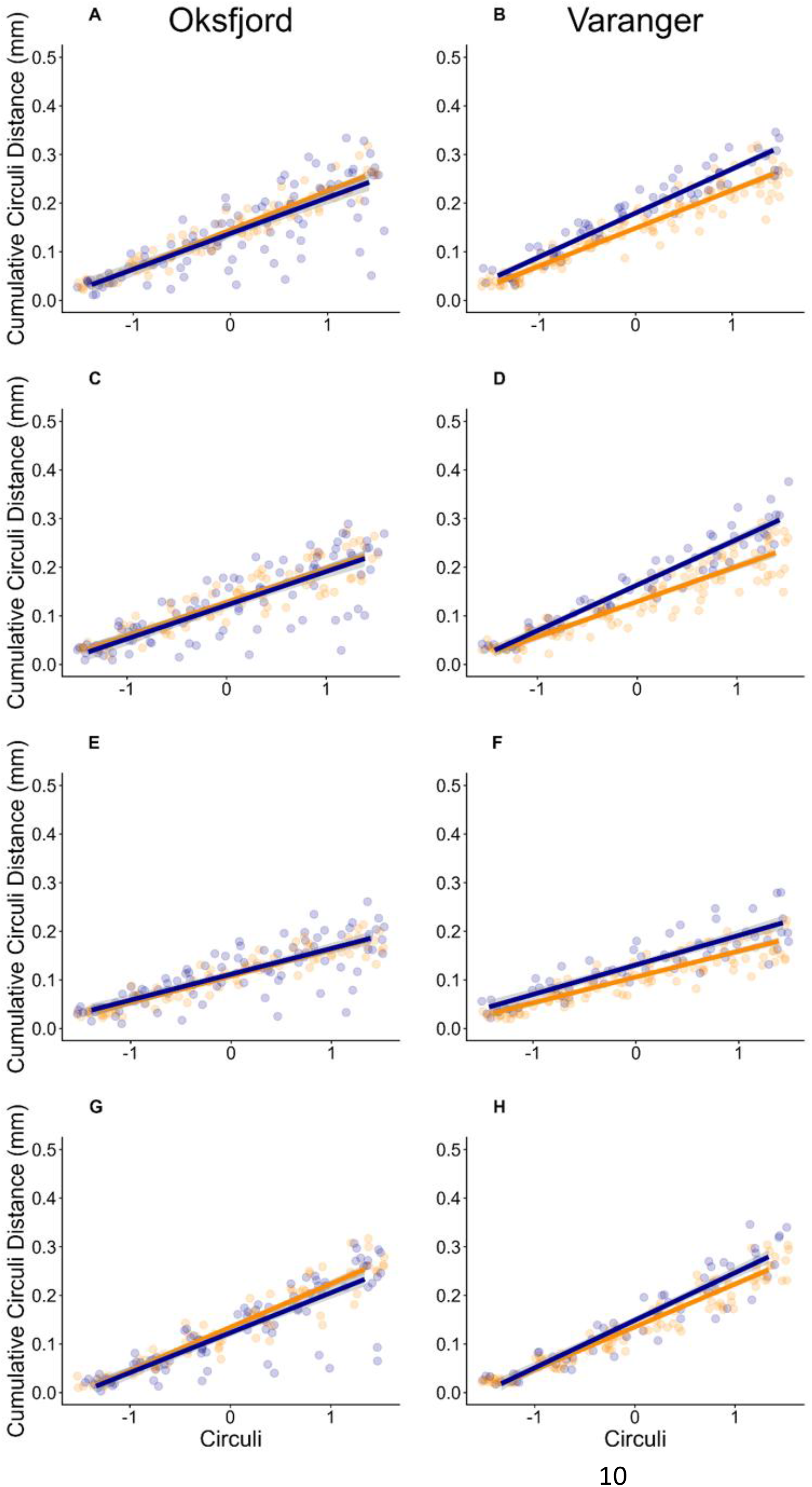
The relationship between cumulative circuli distance (mm) and normalized circuli number (Circuli) for sampling locations for each sex (males in blue, females in orange) within growth periods. A) Period 1, Oksfjord, B) Period 1, Varanger, C) Period 2, Oksfjord, D) Period 2, Varanger, E) Period 3, Oksfjord, F) Period 3, Varanger, G) Period 4, Oksfjord, H) Period 4, Varanger. Colored lines represent linear models, and 95% confidence intervals are represented by grey shading. Small circles show individual data points.

## Discussion

The main objective of this study was to investigate differences in somatic growth of invasive pink salmon caught in two geographically distinct regions of the invasion front in Northern Norway. On average, fishes collected at the two sites during the second summer and at the onset of sexual maturation had similar body lengths, but males were heavier than females. Moreover, males had a significantly higher body condition indicating that they were bulkier than females, likely due to the onset of sexual maturation and the development of secondary sexual traits. Marine growth based on scale analyses varied significantly based on the combination of sex and location. Notably, males from Varanger exhibited a consistently higher growth rate throughout the observed growth periods. These findings suggest that higher early growth was favored in Varanger males, potentially indicating sex-specific selection on faster growth and differences in environmental conditions.

Sexual dimorphism at maturation in pink salmon is well documented in both native and invasive populations and may help to explain why males were heavier and in better condition compared to females. Paulsen et al. (2022) found that males captured in Norway in 2019 and 2021 were significantly longer than females, a pattern consistent with observations from the Northeast Pacific Ocean (Pritchard, 1937). Pink salmon studied in Paulsen et al. (2022) originated from the same odd-year broodline, with a subset collected in rivers with discharge into Varangerfjord in Finnmark, and are thus comparable to the fish investigated in this study. In contrast, we found no significant variation in fish length, but rather weight and condition factor were higher in males than females. One key difference between our study and Paulsen et al. (2022) is that our samples were collected in marine waters prior to pink salmon entering rivers for the return spawning migration and therefore represent an earlier stage of maturation. Moreover, female growth declines during the maturation process as they invest more into gonad development and egg production (Mortensen et al., 2000) while males invest more into secondary sexual traits such as the characteristic hump and kype at the onset of maturation (Davidson, 1935). Thus, the higher weight and condition, but not body length, in males likely reflects early stages of sexual dimorphism in body size and gonadal development related to sexual maturation on their return migration during the summer months.

No significant differences across sex and location were found in either the position of the last winter circulus, or regarding variation in the total amount of circuli deposited. While both circuli spacing, number of circuli, and scale radius reflect somatic growth in pacific salmonids (Fukuwaka & Kaeriyama, 1997), the rate of circuli formation is not fixed. Instead, growth rate is shown to have a positive relationship with circuli formation rate in salmonids (Fukuwaka & Kaeriyama, 1997; Peterson et al., 2021). In pink salmon, formation of circuli has been reported at rates of one circulus every 4.4 days (Pearson, 1966), to one for every 7.4 days (Courtney et al., 2000). Males caught in Varanger showed increased growth rates in this study and it is notable that this is not reflected in either total circuli or the position of the last winter circulus. Given the relatively consistent total number of circuli deposited until the last winter circulus, it can be interpreted that the average circuli formation rate was uniform across the two locations and between the sexes.

All pink salmon in the study followed a pattern of increased growth in the spring/summer followed by limited growth during summer/fall with the lowest growth encountered during fall/winter of year one. Similarly, a substantial increase in growth was experienced during the spring of year two. The pattern of high initial growth tapering off toward autumn is congruent with rapid growth strongly influencing survival during this critical period of early development (Beamish & Mahnken, 2001; Moss et al., 2005). A decline in growth during summer/fall has also been reported in native pink salmon and Atlantic salmon (Mortensen et al., 2000; Todd et al., 2008) and Norwegian invasive pink salmon (Paulsen et al., 2022; Simmons et al., 2026), where growth is likely limited by environmental conditions such as low temperature and prey availability during the onset of autumn (Mortensen et al., 2000). During the fall/winter period, growth declined significantly, a pattern commonly seen in salmonids (Härkönen et al., 2021; Paulsen et al., 2022; Simmons et al., 2026; Todd et al., 2014) and is likely caused by seasonal reductions in temperature, daylight, and food availability.

A variety of factors influence growth in fishes including environmental conditions, density and growth during development and early life, parental effects and the underlying genetic background (Almeida et al., 2024; Maamela et al., 2026; Mobley et al., 2020). The increased sex-specific growth seen in Varanger males raises the question as to whether regional conditions experienced by the pink salmon caught in Varanger are particularly favorable to male growth. The differences observed in growth patterns also indicate that pink salmon returning to Varangerfjord and Oksfjord potentially utilize different marine feeding grounds where they experience different regional conditions. Fish in Oksfjord were sampled 12-17 days later than Varanger, which would theoretically allow for more time for growth and a later maturation stage. Yet, higher growth was seen in Varanger males suggesting that stage of maturation alone cannot explain these differences. While it cannot be concluded directly from this study, environmental factors such as temperature, prey availability or parasitism could disproportionately affect pink salmon in this region. However, data on sex-specific marine migration, growth and size of pink salmon in different regions of the northern Atlantic invasion are limited, and further studies could help shed more light on this relationship.

Size-selective mortality on slow growing Varanger males offers an alternative and non-exclusive explanation to environmental factors to explain higher growth among these individuals. For example, larger, faster-growing juveniles may have a higher survival rate (Moss et al., 2005; Simmons et al., 2026). Because our study only investigates pink salmon that survived to the second summer, factors that may affect mortality prior to this

## Supporting information

Nes et al. Supplemental information

## Acknowledgments

We thank Gunnel Østborg for help with scale imaging, Malte Willmes for valuable guidance on scale analysis, Rachel Paterson for help with scale collection, Eirik Frøiland and the Norwegian Miljø-direktoratet for guidance on pink salmon distributions and the Institute of Marine Research, Tromsø for collections of pink salmon. time are unknown. Thus, it is possible that size-selective mortality on slow growing males is higher in Varanger than in Oksfjord although survivorship bias of the individuals analyzed in this study may mask selection episodes prior to the time of capture.

Numerous factors make it difficult to draw decisive conclusions about pink salmon’s further invasive success in Norway. While scale readings provide a retrospective look into marine growth, the origin and migratory patterns of the fishes in this study are unknown. Future studies that investigate otolith chemistry and kinship analyses may provide indications of the establishment westward along the Norwegian coast (Willmes et al., 2026). Additionally, genetic diversity of pink salmon within Norway appears to be low (Henriksen et al., 2026; Maduna et al., 2024), suggesting that marine growth is unlikely to be the result of local adaptation given the recent and rapid invasion. This low genetic diversity makes invasive pink salmon in Norway an interesting model to study the impacts of environmental factors on their behavior and invasion success.

Understanding how growth differs between life history stages, sexes and locations in invasive species is a fundamental goal of conservation biology. This study reveals a strong pattern of sex-specific somatic growth between two geographic regions along an invasion front. The causes of such a pattern are currently unknown but are likely linked to sexual maturation, environmental conditions and size-selective mortality experienced in pink salmon between the two regions. Due to the scope of this invasion, and its potential to affect native ecosystems, we recommend that future studies should investigate sex-specific growth patterns in more locations and over multiple years to better understand the establishment of the species.

## Author contributions

E.A.N., E.B.T. and K.B.M conceived the study. A.G., K.D., and C.S. and coordinated and/or participated in sample collection. E.A.N. and K.B.M. analyzed the data. E.A.N. and K.B.M drafted the manuscript, with input from all other authors.

## Data and materials availability

All data and code will be available for download at the time of publication.

## Supplementary data

Supplemental data Pink salmon.pdf

## Citations

Almeida, L. Z., Ludsin, S. A., Faust, M. D., & Marschall,E. A. (2024). Lingering legacies: Past growth and parental experience influence somatic growth in a fish population. Journal of Animal Ecology, 93(10), 1462–1474.10.1111/1365-2656.14143

Beacham, T. D., Withler, R. E., & Gould, A. P. (1985). Biochemical Genetic Stock Identification of Pink Salmon (Oncorhynchus gorbuscha) in Southern British Columbia and Puget Sound. Canadian Journal of Fisheries and Aquatic Sciences, 42(9), 1474–1483. 10.1139/f85-185

Beacham, T. D., Withler, R. E., Murray, C. B., & Barner,L. W. (1988). Variation in body size, morphology, egg size, and biochemical genetics of pink salmon in British Columbia. Transactions of the American Fisheries Society, 117(2), 109–126.10.1577/1548-8659(1988)117<0109:VIBSME>2.3.CO;2

Beamish, R. J., & Mahnken, C. (2001). A critical size and period hypothesis to explain natural regulation of salmon abundance and the linkage to climate and climate change. Progress in Oceanography, 49(1), 423–437.10.1016/S0079-6611(01)00034-9

Beamish, R. J., Noakes, D. J., McFarlane, G. A.,Klyashtorin, L., Ivanov, V. V., & Kurashov, V. (1999). The regime concept and natural trends in the production of Pacific salmon. Canadian Journal of Fisheries and Aquatic Sciences, 56(3), 516–526. 10.1139/f98-200

Bjerknes, V. (1977). Evidence of natural production of pink salmon fry (Oncorhynchus gorbuschaWalbaum) in Finnmark, North Norway. Astarte, 105–6.

Brenner, R. E., Moffitt, S. D., & Grant, W. S. (2012). Straying of hatchery salmon in Prince William Sound, Alaska. Environmental Biology of Fishes, 94(1), 179–195. 10.1007/s10641-012-9975-7

Burton, O. J., Phillips, B. L., & Travis, J. M. J. (2010).Trade-offs and the evolution of life-histories during range expansion. Ecology Letters, 13(10), 1210–1220.10.1111/j.1461-0248.2010.01505.x

Christensen, K. A., Rondeau, E. B., Sakhrani, D., Biagi,C. ., Johnson, H., Joshi, J., Flores, A.-M.,Leelakumari, S., Moore, R., Pandoh, P. K., Withler,R. E., Beacham, T. D., Leggatt, R. A., Tarpey, C. M., Seeb, L. W., Seeb, J. E., Jones, S. J. M., Devlin, R. H., & Koop, B. F. (2021). The pink salmon genome: Uncovering the genomic consequences of a two-year life cycle. PLoS One, 16(12), e0255752.10.1371/journal.pone.0255752

Courtney, D. L., Mortensen, D. G., & Orsi, J. A. (2000).Digitized scale and otolith microstructures as correlates of juvenile pink salmon size. North Pacific Anadromous Fish Commission Bulletin, 2, 337–345.

Davidson, F. A. (1935). The development of the secondary sexual characters in the pink salmon (Oncorhynchus gorbuscha). Journal of Morphology, 57(1), 169–183.10.1002/jmor.1050570110

Diaz Pauli, B., Berntsen, H. H., Thorstad, E. B., Homrum, E. í., Lusseau, S. M., Wennevik, V., & Utne, K. R. (2022). Geographic distribution, abundance, diet, and body size of invasive pink salmon (Oncorhynchus gorbuscha) in theNorwegian and Barents Seas, and in Norwegian rivers. ICES Journal of Marine Science, 80(1), 76–90. 10.1093/icesjms/fsac224

Dickerson, B. R., Quinn, T. P., & Willson, M. F. (2002).Body size, arrival date, and reproductive success of pink salmon, Oncorhynchus gorbuscha. Ethology Ecology & Evolution, 14(1), 29–44.10.1080/08927014.2002.9522759

Dunlop, K., Eloranta, A. P., Gjelland, K. Ø., Slåteng, S.,Jensen, J., Knudsen, R., Kiljunen, M., Jensen, H., Svenning, M. A., Muladal, R., & Frainer, A. (2026). The feeding ecology of invasive pink salmonjuveniles in northern Norwegian rivers and their role as prey to native salmonids. Ecology Of Freshwater Fish, 35(3), e70047.10.1111/eff.70047

Erkinaro, J., Orell, P., Fossøy, F., Kytökorpi, M., Gjelland,K., Johansen, N., Domaas, S., Kuusela, J., Fagard, P., Frøiland, E., & Falkegård, M. (2026). Rapid increase in abundance and distribution of invasive pink salmon (Oncorhynchus gorbuscha) within adiverse, large Barents Sea catchment. Journal of Fish Biology, 108(6), 1875–1888.10.1111/jfb.70348

Erkinaro, J., Orell, P., Kytökorpi, M., Pohjola, J.-P., & Power, M. (2024). Active feeding of downstream migrating juvenile pink salmon (Oncorhynchus gorbuscha) revealed in a large Barents Sea river using diet and stable isotope analysis. Journal of Fish Biology, 104(3), 797–806. 10.1111/jfb.15625

Fukuwaka, M. (1998). Scale and otolith patterns provegrowth history of Pacific salmon. North PacificAnadromous Fish Commission Bulletin, 1, 190–198.

Fukuwaka, M., & Kaeriyama, M. (1997). Scale analyses to estimate somatic growth in sockeye salmon, Oncorhynchus nerka. Canadian Journal of Fisheries and Aquatic Sciences, 54(3), 631–636.10.1139/f96-307

Gallagher, Z. S., Bystriansky, J. S., Farrell, A. P., & Brauner, C. J. (2013). A novel pattern ofsmoltification in the most anadromous salmonid: pink salmon (Oncorhynchus gorbuscha). Canadian Journal of Fisheries and Aquatic Sciences, 70(3), 349–357. 10.1139/cjfas-2012-0390

Gende, S. M., Quinn, T. P., Willson, M. F., Heintz, R., & Scott, T. M. (2004). Magnitude and fate of salmon-derived nutrients and energy in a coastal stream ecosystem. Journal of Freshwater Ecology, 19(1), 149–160.10.1080/02705060.2004.9664522

Gordeeva, N. V., Salmenkova, E. A., & Prusov, S. V.(2015). Variability of biological and population genetic indices in pink salmon, Oncorhynchus gorbuscha transplanted into the White Sea basin. Journal of Ichthyology, 55(1), 69–76.10.1134/S0032945215010051

Hartig, F. (2026). DHARMa: Residual Diagnostics forHierarchical (Multi-Level / Mixed) Regression Models. In

Hawkins, S. L., Varnavskaya, N. V., Matzak, E. A., Efremov, V. V., Guthrie III, C. M., Wilmot, R. L.,Mayama, H., Yamazaki, F., & Gharrett, A. J. (2002). Population structure of odd-broodline Asian pink salmon and its contrast to the even-broodlinestructure. Journal of Fish Biology, 60(2), 370–388. 10.1111/j.1095-8649.2002.tb00287.x

Heard, W. R. (1991). Life history of pink salmon (Oncorhynchus gorbuscha). In C. Groot & L. Margolis (Eds.), Pacific Salmon Life Histories.UBC Press.

Heincke, F. (1908). Bericht über dieUntersuchungen der Biologischen Anstaltauf Helgoland zurNaturgeschichte der Nutzfische. Die Beteiligung Deutschlandsan der InternationalenMeeresforschung, 1908:4(5), 67–155.

Hendry, A. P., Castric, V., Kinnison, M. T., & Quinn, T. P. (2003). The evolution of philopatry and dispersal: homing versus straying in salmonids. In A. P. Hendry & S. C. Sterns (Eds.), Evolution Illuminated: Salmon and Their Relatives (pp. 53–91). Oxford University Press.

Henriksen, E. H., Maduna, S. N., Hagen, S. B., & Christensen, G. N. (2026). Genomic evidence of mainland-driven invasion without localestablishment of pink salmon in Svalbard, Norway. Scientific Reports. 10.1038/s41598-026-69339-z

Hindar, K., Hole, L. R., Kausrud, K., Malmstrøm, M., Rimstad, E., Robertson, L., Sandlund, O. T.,Thorstad, E. B., Vollset, K. W., de Boer, H., Eldegard, K., Järnegren, J., Kirkendall, L., Måren, I., Nielsen, A., Nilsen, E. B., Rueness, E., & Velle, G. (2020).Assessment of the risk to Norwegian biodiversity and aquaculture from pink salmon (Oncorhynchus gorbuscha). VKM report 2020:01 (ScientificOpinion of the Panel on Alien Organisms and Trade in Endangered Species (CITES) of the Norwegian Scientific Committee for Food and Environment,Issue. https://hdl.handle.net/11250/2729831

Härkönen, L., Louhi, P., Huusko, R., & Huusko, A.(2021). Wintertime growth in Atlantic salmon under changing climate: the importance of ice cover for individual growth dynamics. Canadian Journal of Fisheries and Aquatic Sciences, 78(10), 1479–1485. 10.1139/cjfas-2020-0258

Iritani, A. T., Barnes, E. M., & Phelps, M. P. (2025).Influence of egg size and parental genetics on the metabolic rate of Chinook and pink salmonembryos. Conserv Physiol, 13(1), coaf062. 10.1093/conphys/coaf062

Jeffries, K. M., Hinch, S. G., Martins, E. G., Clark, T. D., Lotto, A. G., Patterson, D. A., Cooke, S. J., Farrell, A. P., & Miller, K. M. (2011). Sex and proximity to reproductive maturity influence the survival, final maturation, and blood physiology of Pacific salmon shen exposed to high temperature during asimulated migration. Physiological and Biochemical Zoology, 85, 62–73.

Kaev, A. M. (2015). Retrospective analysis of growth of pink salmon Oncorhynchus gorbuscha from scales: Interannual variability. Journal of Ichthyology, 55(5), 707–722.10.1134/S0032945215050070

Kaev, A. M., Romasenko, L. V., & Kaev, D. A. (2023).Scale-based study of linear growth rate of pink salmon Oncorhynchus gorbuscha (Salmonidae). Journal of Ichthyology, 63(1), 79–89.10.1134/S0032945223010058

Kovach, R. P., Gharrett, A. J., & Tallmon, D. A. (2012).Genetic change for earlier migration timing in a pink salmon population. Proceedings of the Royal Society B: Biological Sciences, 279(1743), 3870–3878. 10.1098/rspb.2012.1158

Kuznetsova, A., Brockhoff, P. B., & Christensen, R. H. B.(2017). lmerTest Package: tests in Linear Mixed Effects Models Journal of Statistical Software, 82(13), 1–26.10.18637/jss.v082.i13

Kwain, W., & Rose, G. A. (1986). Spawning migration of Great Lakes pink salmon (Oncorhynchus Gorbuscha): size and sex distributions, riverentrance and exit. Journal of Great Lakes Research, 12(2), 101–108.10.1016/S0380-1330(86)71704-8

Langan, J. A., Alsip, P. J., Abdelhady, H. U., Bronte, C. R., Goldsworthy, C. A., Kornis, M. S., Oke, K. B., Thorstad, E. B., & Turschak, B. A. (2026). The forgotten pink salmon in the Laurentian Great Lakes: an unexpected invasion with insights forthree oceans. Fish and Fisheries, 27(4), 813–830. 10.1111/faf.70084

Lennox, R. J., Berntsen, H. H., Garseth, Å. H., Hinch, S. G., Hindar, K., Ugedal, O., Utne, K. R., Vollset, K. W., Whoriskey, F. G., & Thorstad, E. B. (2023).Prospects for the future of pink salmon in three oceans: From the native Pacific to the novel Arcticand Atlantic. Fish and Fisheries, 24(5), 759–776. 10.1111/faf.12760

Maamela, K. S., Prokkola, J. M., Suvanto, C., Huang, X.-D., Primmer, C. R., & Mobley, K. B. (2026). Theinfluence of parental and genotype effects on early survival and development in Atlantic salmon.bioRxiv, 2026.2008.2014.744584.10.64898/2026.08.14.744584

Maduna, S. N., Aspholm, P. E., Hansen, A.-S. B.,Klütsch, C. F. C., & Hagen, S. B. (2024). Ecological niche modelling and population genomics provide insights into the geographic and demographic‘explosion’ of a non-indigenous salmonid. Diversity and Distributions, 30(3), e13811.10.1111/ddi.13811

Miljø-direktoratet. (2026). Pukkellaks-uttak i 2023.Retrieved 27.10.2026 from https://www.miljodirektoratet.no/aktuelt/datavisualisering/pukkellaks-uttak-2023/

Mo, T. A., Thorstad, E. B., Sandlund, O. T., Berntsen, H.H., Fiske, P., & Uglem, I. (2018). The pink salmoninvasion: a Norwegian perspective. Journal of Fish Biology, 93(1), 5–7.10.1111/jfb.13682

Mobley, K. B., Granroth-Wilding, H., Ellmen, M., Orell, P., Erkinaro, J., & Primmer, C. R. (2020). Time spent in distinct life-history stages has sex-specific effects on reproductive fitness in wild Atlantic salmon.Molecular Ecology, 29(6), 1173–1184. 10.1111/mec.15390

Mortensen, D., Wertheimer, A., Taylor, S., & Landingham, J. (2000). The relation between early marine growth of pink salmon, Oncorhynchus gorbuscha, and marine water temperature, secondary production, and survival to adulthood. Fisheries Bulletin, 98, 319–335.

Mortensen, D. G., Wertheimer, A. C., Maselko, J. M., & Taylor, S. G. (2002). Survival and straying of Auke Creek, Alaska, pink salmon marked with codedwire tags and thermally induced otolith marks. Transactions of the American Fisheries Society, 131(1), 14–26. 10.1577/1548-8659(2002)131<0014:Sasoac>2.0.Co;2

Moss, J. H., Beauchamp, D. A., Cross, A. D., Myers, K.W., Farley, E. V., Jr., Murphy, J. M., & Helle, J. H. (2005). Evidence for size-selective mortality after the first summer of ocean growth by pink salmon. Transactions of the American Fisheries Society, 134(5), 1313–1322. 10.1577/t05-054.1

Paulsen, T., Sandlund, O. T., Østborg, G., Thorstad, E. B., Fiske, P., Muladal, R., & Tronstad, S. (2022).Growth of invasive pink salmon (Oncorhynchus gorbuscha) at sea assessed by scale analysis. Journal of Fish Biology, 100(1), 218–228.10.1111/jfb.14937

Pearson, R. E. (1966). Number of circuli and time of annulus formation on scales of pink salmon (Oncorhynchus gorbuscha). Journal of the Fisheries Research Board of Canada, 23(5), 747–756.10.1139/f66-065

Perkins, T. A., Phillips, B. L., Baskett, M. L., & Hastings,A. (2013). Evolution of dispersal and life historyinteract to drive accelerating spread of an invasive species. Ecology Letters, 16(8), 1079–1087.10.1111/ele.12136

Pess, G. R., Hilborn, R., Kloehn, K., & Quinn, T. P.(2012). The influence of population dynamics and environmental conditions on pink salmon (Oncorhynchus gorbuscha) recolonization after barrier removal in the Fraser River, British Columbia, Canada. Canadian Journal of Fisheries and Aquatic Sciences, 69(5), 970–982. 10.1139/f2012-030

Peterson, E., Sheehan, T. F., & Zydlewski, J. (2021). Scale growth rates and scale circulus deposition rates of marine-stage Atlantic salmon Salmo salar raised under semi-natural conditions. Journal of Northwest Atlantic Fishery Science, 52, 19–27. 10.2960/J.v52.m733

Phillips, B. L., & Shine, R. (2006). An invasive speciesinduces rapid adaptive change in a native predator: cane toads and black snakes in Australia.Proceedings of the Royal Society B-Biological Sciences, 273(1593), 1545–1550.10.1098/rspb.2006.3479

Pritchard, A. L. (1937). Variation in the time of run, sex proportions, size and egg content of adult pink salmon (Oncorhynchus gorbuscha) at McClinton Creek, Masset Inlet, B.C. Journal of the BiologicalBoard of Canada, 3(5), 403–416. 10.1139/f37-023

Pyper, B. J., Mueter, F. J., Peterman, R. M., Blackbourn,D. J., & Wood, C. C. (2001). Spatial covariation in survival rates of Northeast Pacific pink salmon (Oncorhynchus gorbuscha). Canadian Journal of Fisheries and Aquatic Sciences, 58(8), 1501–1515. 10.1139/f01-096

Quinn, T. P. (2018). The Behavior and Ecology of Pacific Salmon and Trout (2nd ed.). University of Washington Press.

R Core Team. (2026). R: A Language and Environment for Statistical Computing. In R Foundation for Statistical Computing.

Sakai, A. K., Allendorf, F. W., Holt, J. S., Lodge, D. M.,Molofsky, J., With, K. A., Baughman, S., Cabin, R. J., Cohen, J. E., Ellstrand, N. C., McCauley, D. E., apos, Neil P., Parker, I. M., Thompson, J. N., & Weller, S.G. (2001). The population biology of invasive species. Annual Review of Ecology, Evolution, and Systematics, 32(Volume 32, 2001), 305–332.10.1146/annurev.ecolsys.32.081501.114037

Sandlund, O. T., Berntsen, H. H., Fiske, P., Kuusela, J., Muladal, R., Niemelä, E., Uglem, I., Forseth, T., Mo,T. A., Thorstad, E. B., Veselov, A. E., Vollset, K. W., & Zubchenko, A. V. (2019). Pink salmon in Norway:the reluctant invader. Biological Invasions, 21(4), 1033–1054. 10.1007/s10530-018-1904-z

Schneider, C. A., Rasband, W. S., & Eliceiri, K. W. (2012). NIH Image to ImageJ: 25 years of image analysis. Nature Methods, 9(7), 671–675. 10.1038/nmeth.2089

Shine, R. (2012). Invasive species as drivers of evolutionary change: cane toads in tropicalAustralia. Evolutionary Applications, 5(2), 107–116. 10.1111/j.1752-4571.2011.00201.x

Simmons, O. M., Paulsen, T., Berntsen, H. H., Diserud, O., Fiske, P., Østborg, G., Willmes, M., & Thorstad,E. B. (2026). Early marine growth and spawning body size vary by year and region in invasive pink salmon (Oncorhynchus gorbuscha). AquaticSciences, 88(3), 77.10.1007/s00027-026-01309-z

Staveley, T. A. B., Bergendahl, I. A., Bárðarson, H.,Berntsen, H. H., Eliasen, K., Erkinaro, J., Nygaard, R., Sivebæk, F., & Thorstad, E. B. (2025). Status and future perspectives of pink salmon in the Nordic region. Boreal Environment Research, 30, 149–162. 10.60910/ber2025.wt02-y377

Thorstad, E. B., Frøiland, E., Fagard, P., Kvaal, M., Lehne, C. K., Liberg, E., Sandodden, R., Skaala, Ø., & Vatne, T. (2025). Assessing the extraordinary measures used to counteract invasion of pink salmon into the North Atlantic. Management ofBiological Invasions, 16, 465–487.10.3391/mbi.2025.16.2.09

Thorstad, E. B., Frøiland, E., Fagard, P., Lehne, C. K., Liberg, E., Muladal, R., Østborg, G., Sandodden, R.,Vatne, T., & Vistnes, H. (2026). Status and prospects of introduced even-year pink salmon inthe North Atlantic and Barents Sea regions. Boreal Environment Research, 31, 63–74.10.60910/ber2026.peam-tn54

Todd, C. D., Hughes, S. L., Marshall, C. T., MacLean, J.C., Lonergan, M. E., & Biuw, E. M. (2008).Detrimental effects of recent ocean surface warming on growth condition of Atlantic salmon. Global Change Biology, 14(5), 958–970.10.1111/j.1365-2486.2007.01522.x

Todd, C. D., Whyte, B. D. M., MacLean, J. C., Revie, C.W., Lonergan, M. E., & Hanson, N. N. (2014). A simple method of dating marine growth circuli on scales of wild one sea-winter and two sea-winter Atlantic salmon (Salmo salar). Canadian Journal of Fisheries and Aquatic Sciences, 71(5), 645–655. 10.1139/cjfas-2013-0359

Walker, B. M., & Sutton, T. M. (2016). Growth-increment formation using otoliths and scales for age-0 chinook salmon. North American Journal of Fisheries Management, 36(5), 995–999.10.1080/02755947.2016.1184202

Willmes, M., Wacker, S., Berntsen, H. H., Lee, E.,Løkeberg, G., Huyskens, M. H., James, H. F., Snoeck, C., Thorstad, E. B., & Karlsson, S. (2026). Combining kinship analysis and otolith geochemistry toinvestigate homing and straying of invasive pink salmon (Oncorhynchus gorbuscha). Boreal Env. Res., 31, 91–100.10.60910/ber2026.9mgy-5v85

