## Supplementary material for "Sex-specific growth in invasive pink salmon (*Oncorhynchus gorbuscha*) from two regions in northern Norway": Nes et al. Supplemental information

This file includes:

Tables S1-S3

Figure S1

Table S1. Summary of body size estimates of sampled pink salmon by location and sex, including sample sizes (n) and the mean for both sexes (Total). Means (±SE) are given for measured fork length (cm), weight (kg), and condition factor (*K*).

| **Location** | **Sex** | ***n*** | **Fork length (cm)** | **Weight (kg)** | ***K*** |
|---|---|---|---|---|---|
| Oksfjord | Female | 14 | 45.6 ± 0.9 | 1.36 ± 0.07 | 1.42 ± 0.03 |
| | Male | 16 | 45.6 ± 1.4 | 1.49 ± 0.12 | 1.53 ± 0.04 |
| | Total | 30 | 45.6 ± 0.8 | 1.43 ± 0.07 | 1.48 ± 0.02 |
| Varanger | Female | 19 | 45.4 ± 0.5 | 1.41 ± 0.04 | 1.50 ± 0.03 |
| | Male | 11 | 48.1 ± 0.9 | 1.81 ± 0.12 | 1.61 ± 0.07 |
| | Total | 30 | 46.4 ± 0.5 | 1.56 ± 0.06 | 1.54 ± 0.03 |

Table S2. Summary of scale measurements of sampled pink salmon by location and sex, including sample sizes (*n*) and the mean for both sexes (Total). Means (±SE) are given for total number of circuli (total circuli), position of the last winter circulus counted from the focus (LWC position), distance from focus to last winter circulus (LWC distance, mm), and mean distance between circuli (Circuli distance, mm).

| **Location** | **Sex** | ***n*** | **Total circuli** | **LWC position** | **LWC distance (mm)** | **Circuli distance (mm)** |
|---|---|---|---|---|---|---|
| Oksfjord | Female | 14 | 30.9 ± 0.5 | 19.6 ± 0.3 | 0.76 ± 0.02 | 0.038 ± 0.001 |
| | Male | 16 | 30.5 ± 0.7 | 19.5 ± 0.7 | 0.70 ± 0.06 | 0.035 ± 0.002 |
| | Total | 30 | 30.6 ± 0.5 | 19.5 ± 0.4 | 0.73 ± 0.03 | 0.037 ± 0.001 |
| Varanger | Female | 19 | 30.9 ± 0.5 | 19.5 ± 0.4 | 0.76 ± 0.03 | 0.039 ± 0.001 |
| | Male | 10 | 32.0 ± 0.8 | 21.0 ± 0.6 | 0.91 ± 0.03 | 0.044 ± 0.001 |
| | Total | 29 | 31.3 ± 0.4 | 20.0 ± 0.3 | 0.81 ± 0.02 | 0.040 ± 0.001 |

Table 3. Results of general linear models showing the effects of non-normalized circuli number (circuli no.), sex and location on cumulative circuli distance (mm) for growth periods 1-4. Reference sex is female, reference location is Oxfjord. Significance for bold values indicates α = 0.05.

| Response variable: Cumulative circuli distance | | | | | |
|---|---|---|---|---|---|
| | **Predictor** | **Estimate** | **SE** | **t** | **Pr(>\|t\|)** |
| Period 1 Spring/Summer Year 1 | Intercept | -0.0414 | 0.0079 | -5.266 | **<0.0001** |
| | Circuli no. | 0.0388 | 0.0008 | 49.388 | **<0.0001** |
| | Sex | 0.0056 | 0.0107 | 0.517 | 0.6063 |
| | Location | 0.0078 | 0.0104 | 0.755 | 0.4521 |
| | Circuli no.*Sex | -0.0032 | 0.0011 | -3.001 | **0.0029** |
| | Circuli no.*Location | -0.0009 | 0.0010 | -0.897 | 0.3702 |
| | Sex*Location | -0.0015 | 0.0157 | -0.096 | 0.9241 |
| | Circuli no.*Sex*Location | 0.0070 | 0.0015 | 4.622 | **<0.0001** |
| Period 2 Summer/Fall Year 1 | Intercept | -0.2708 | 0.0128 | -21.187 | **<0.0001** |
| | Circuli no. | 0.0357 | 0.0009 | 40.198 | **<0.0001** |
| | Sex | -0.0051 | 0.0174 | -0.295 | 0.7682 |
| | Location | -0.0148 | 0.0168 | -0.882 | 0.3789 |
| | Circuli no.*Sex | -0.0004 | 0.0012 | -0.319 | 0.7498 |
| | Circuli no.*Location | 0.0014 | 0.0012 | 1.226 | 0.2210 |
| | Sex*Location | -0.0789 | 0.0251 | -3.143 | **0.0019** |
| | Circuli no.*Sex*Location | 0.0081 | 0.0017 | 4.802 | **<0.0001** |
| Period 3: Fall/winter Year 1 | Intercept | -0.3993 | 0.0170 | -23.513 | **<0.0001** |
| | Circuli no. | 0.0296 | 0.0008 | 37.862 | **<0.0001** |
| | Sex | 0.0238 | 0.0231 | 1.031 | 0.3034 |
| | Location | 0.0095 | 0.0224 | 0.425 | 0.6714 |
| | Circuli no.*Sex | -0.0013 | 0.0011 | -1.198 | 0.2317 |
| | Circuli no.*Location | -0.0007 | 0.0010 | -0.656 | 0.5124 |
| | Sex*Location | -0.0671 | 0.0335 | -2.004 | **0.0462** |
| | Circuli no.*Sex*Location | 0.0030 | 0.0015 | 1.979 | **0.0488** |
| Period 4: Spring/Summer Year 2 | Intercept | -0.0324 | 0.0079 | -4.087 | **<0.0001** |
| | Circuli no | 0.0477 | 0.0011 | 45.920 | **<0.0001** |
| | Sex | 0.0018 | 0.0109 | 0.164 | 0.8702 |
| | Location | 0.0029 | 0.0105 | 0.273 | 0.7802 |
| | Circuli no.*Sex | -0.0037 | 0.0015 | -2.458 | **0.0145** |
| | Circuli no.*Location | -0.0006 | 0.0015 | -0.428 | 0.6688 |
| | Sex*Location | -0.0061 | 0.0159 | -0.381 | 0.7044 |
| | Circuli no.*Sex*Location | 0.0089 | 0.0022 | 3.986 | **<0.0001** |

Figure S1. The relationship between cumulative circuli distance (mm) and circuli number (Circuli No.) for Oksfjord and Varanger sampling locations for each sex (Males in blue, females in orange) within growth periods. A) Period 1, Oksfjord, B) Period 1, Varanger, C) Period 2, Oksfjord, D) Period 2, Varanger, E) Period 3, Oksfjord, F) Period 3, Varanger, G) Period 4, Oksfjord, H) Period 4, Varanger. Cumulative distance represents the cumulative growth from the previous circuli. Colored lines represent linear models, and 95% confidence intervals are represented by grey shading. Colored dots represent individual data points.

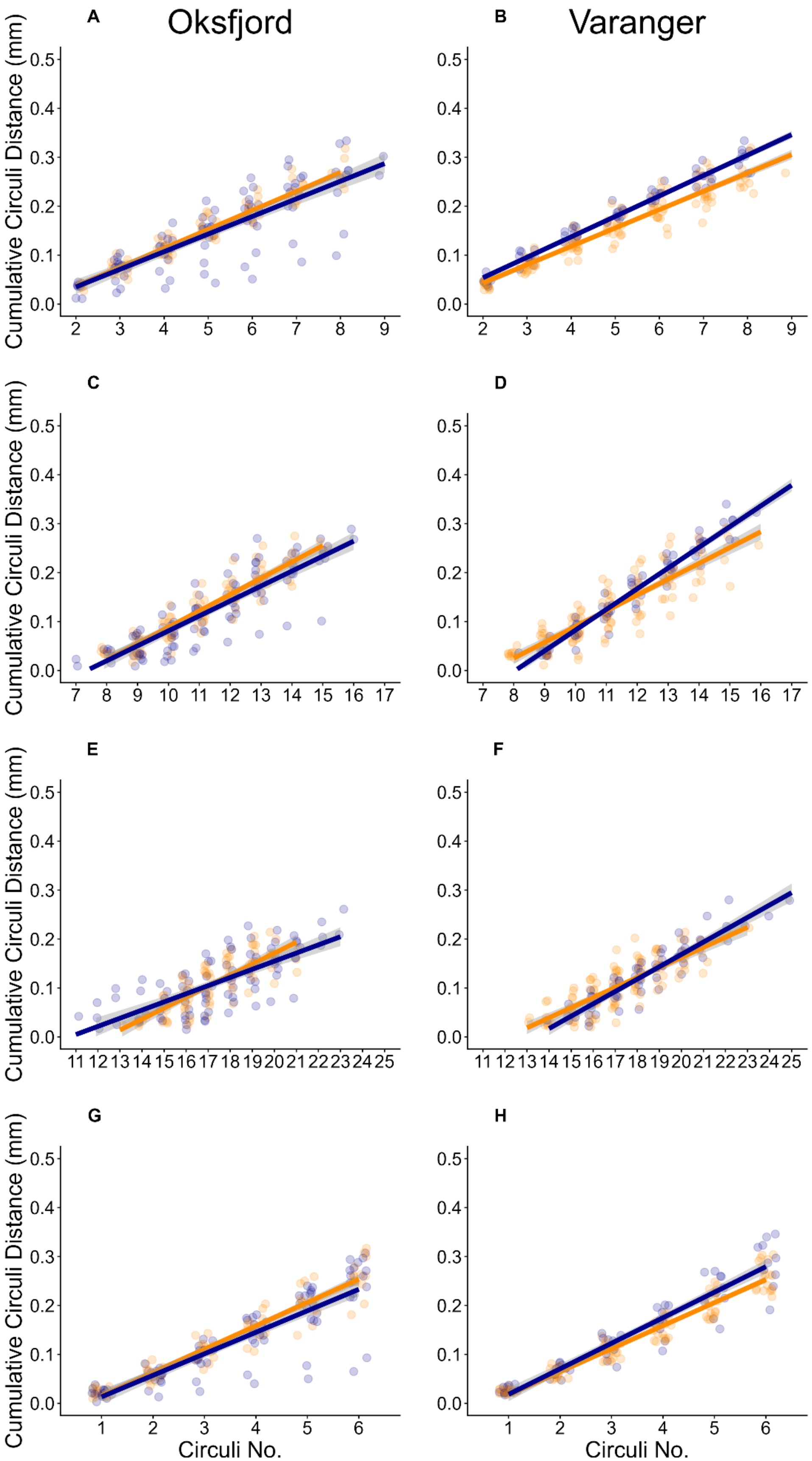
A
Oksfjord
B
Varanger
C
D
E
F
G
H
Cumulative Circuli Distance (mm)
Circuli No.
